# The genome of the coffee bean weevil (*Araecerus fasciculatus*) reveals a cytochrome P450 repertoire as a convergent candidate mechanism of insect-origin caffeine detoxification

**DOI:** 10.64898/2026.06.11.731724

**Authors:** Laura V. Martínez Aponte, Alfredo Rodriguez Ruiz, Sean A. Locke, Timothy J. Colston, Alex R. Van Dam

**Affiliations:** Department of Biology, University of Puerto Rico, Mayagüez, PR 00681, USA; Department of Biological Sciences, Texas Tech University; Genomic Resources Collection, University of Puerto Rico, Mayagüez, PR 00681, USA; Museum für Naturkunde Berlin, Leibniz-Institut für Evolutions- und Biodiversitätsforschung, 10115 Berlin, Germany

**Keywords:** *Araecerus fasciculatus*, Anthribidae, genome assembly, cytochrome P450, caffeine detoxification, horizontal gene transfer, PacBio HiFi, coffee pest, Curculionoidea

## Abstract

The coffee bean weevil, *Araecerus fasciculatus* (Coleoptera, Curculionoidea, Anthribidae), is a cosmopolitan pest of over 100 stored agricultural commodities, with particular economic impact on coffee (*Coffea arabica*). Although two chromosome-level anthribid genomes have recently been released as part of the Darwin Tree of Life (DToL) project (Booth et al. 2024; Crowley et al. 2025), no functionally annotated genome has been available for the family. Here we present a draft genome assembly for *A. fasciculatus*, generated from PacBio HiFi long reads and processed through a three tiered metagenomic filtering pipeline to remove host plant (*C. arabica*) and microbial contamination. The final assembly spans 475 Mb across 3,617 scaffolds (N50 = 170 kb) with 88.5% BUSCO completeness (insecta_odb10) and only 3.1% duplication. Gene prediction with BRAKER2 identified 22,384 protein-coding genes, of which 11,783 received functional annotations through SwissProt similarity. Notably, we identified 92 cytochrome P450 (CYP) genes, including tandem gene clusters on two scaffolds (4 genes on ptg000464l, 5 genes on ptg001867l), suggestive of lineage-specific expansion through tandem duplication. Homology searches against *Drosophila melanogaster* caffeine-metabolizing P450s (CYP12D1, CYP6d5, CYP6a8) recovered strong matches (e-values 9.7 × 10^-110^ to 5.4 × 10^-101^, 33–38% identity). In stark contrast, comprehensive BLAST searches for bacterial caffeine N-demethylase genes (ndmA/B/C/D), which mediate caffeine degradation via horizontal gene transfer in the coffee berry borer *Hypothenemus hampei* (Scolytinae), returned zero hits across the *A. fasciculatus* genome, predicted proteome, and associated bacterial scaffolds. AlphaFold2 structure prediction of four top *Araecerus* P450 candidates produced high-confidence models (pLDDT 84.5–93.9, pTM 0.735–0.930) with conserved P450 catalytic motifs. Foldseek structural homology searches confirmed that all four candidates adopt cytochrome P450 folds (top hits: human CYP3A4, CYP3A7, CYP11A1; TM-scores 0.90–0.92; probability 1.000), with zero hits to bacterial Rieske-fold enzymes. Molecular docking of caffeine against these structures yielded binding affinities of −5.41 to −5.80 kcal/mol for the *Araecerus* candidates, comparable to or exceeding the −5.55 kcal/mol obtained for the experimentally validated *Drosophila* CYP6a8 and substantially stronger than the −3.70 kcal/mol for the bacterial NdmA structural outgroup (PDB: 6ICP). Phylogenetic analysis revealed that all four candidates have clear orthologs in two non-seed-feeding DToL anthribids (*Pseudeuparius sepicola* and *Platystomos albinus*), demonstrating that these P450 genes predate the dietary transition to caffeine-containing seeds. The *Araecerus* candidates predominantly belong to the CYP6 family (clan 3), whereas the primary *Drosophila* caffeine P450 CYP12D1 belongs to the mitochondrial clan, confirming convergent recruitment of different P450 subfamilies for caffeine metabolism. These results support the hypothesis that *A. fasciculatus* employs an insect-encoded, P450-mediated caffeine detoxification pathway fundamentally distinct from the bacterial horizontal gene transfer mechanism documented in Scolytinae. This represents convergent evolution of caffeine resistance via independent molecular strategies within Curculionoidea, and provides the first functionally annotated genomic resource for comparative studies across the Anthribidae.

## Introduction

*Araecerus fasciculatus* (DeGeer, 1775) (Coleoptera, Curculionoidea, Anthribidae), commonly known as the coffee bean weevil, is among the most economically damaging pests of stored agricultural commodities worldwide. First described as *Curculio fasciculatus* by Charles DeGeer and later placed in *Araecerus* by Schoenherr (1823), the species has been recorded as a pest of over 100 plant products including coffee, cocoa, nutmeg, dried fruits, cassava, and various spices (Alba-Alejandre et al. 2018; Caasi-Lit and Lit 2011). Originally endemic to the Indo-Pacific region, *A. fasciculatus* has achieved a cosmopolitan distribution through the global trade of infested commodities, and is now established across tropical and subtropical regions of the Americas, Africa, Asia, Australia, and Europe (Childers and Woodruff 1980; Valentine 2005).

Coffee (*Coffea arabica* and *C. canephora*) represents one of the most economically significant hosts, with the global coffee trade valued at over $70 billion annually (Vega et al. 2006). Female *A. fasciculatus* oviposit on dried coffee pulp, larvae penetrate and consume the seed endosperm, and pupate inside the fruit (Fornazier et al. 2019). Adults emerge by boring exit holes, damaging both the cherry exterior and the bean, thereby reducing crop yield and market value (Alba-Alejandre et al. 2018; Childers 1982). Despite its economic significance, *A. fasciculatus* has received far less genomic attention than the primary coffee pest *Hypothenemus hampei* (Scolytinae), the coffee berry borer, for which a draft genome has been available since 2015 (Vega et al. 2015).

A central question in coffee beetle biology is how these insects metabolize caffeine (1,3,7-trimethylxanthine), a purine alkaloid that serves as a natural insecticide in coffee plants (Nathanson 1984). In insects, two fundamentally different biochemical mechanisms of caffeine metabolism have been documented. In *Drosophila melanogaster*, caffeine is metabolized through insect-encoded cytochrome P450 monooxygenases, particularly CYP12D1, CYP6d5, and CYP6a8 (Coelho et al. 2015). This represents an intrinsic detoxification capability. In contrast, *H. hampei* appears to rely at least in part on horizontally transferred bacterial genes integrated into its genome, with ten putative horizontal gene transfer (HGT) events documented (Vega et al. 2015). Additionally, caffeine-degrading gut bacteria harboring N-demethylase genes (ndmA, ndmB, ndmC, ndmD) have been isolated from *H. hampei* (Vega et al. 2021). Whether other coffee-feeding beetles employ one or both of these strategies remains unknown.

The Anthribidae (fungus weevils) are phylogenetically distant from the Scolytinae within Curculionoidea, making *A. fasciculatus* an ideal system for testing whether caffeine detoxification is conserved within this diverse beetle superfamily (i.e., employs HGT derived genes as in Scolytinae) or has evolved convergently via different molecular mechanisms, such as using the intrinsic genes found in *Drosophila*. While two Anthribidae genomes have recently been released through the Darwin Tree of Life project (*Pseudeuparius sepicola*, Booth et al. 2024; *Platystomos albinus*, Crowley et al. 2025), these chromosome-level assemblies lack gene prediction and functional annotation. Here we present the first functionally annotated anthribid genome, and use comparative genomics, molecular docking, and protein structure prediction to investigate the molecular basis of caffeine metabolism in *A. fasciculatus*.

## Materials and Methods

### Sample collection and DNA extraction

Adult *A. fasciculatus* specimens were collected from a 60 kg bag of roasted coffee beans at Café Mis Abuelos coffee farm in Mayagüez, Puerto Rico on December 2, 2021. Specimens were maintained in plastic storage containers at the UPRM Invertebrate Collection Laboratory. DNA extractions were performed using three kits: Circulomics Nanobind CBB Big DNA Kit (now PacBio), Qiagen PureGene Tissue Kit, and Qiagen MagAttract High Molecular Weight DNA Kit, for a total of 12 extractions from whole-body specimens homogenized with micropestles. DNA was quantified using a Promega Quantus Fluorometer. The highest-yield sample (AF-12, Qiagen MagAttract, 9.9 ng/μ) was selected for sequencing.

### Sequencing

Sample AF-12 was shipped to the University of Maryland Genomic Resource Center and assessed for quality using an Agilent 5200 Fragment Analyzer System. A PacBio HiFi library was prepared and sequenced on a single SMRT Cell using the PacBio Sequel II platform in February 2022.

### Genome assembly and contamination filtering

CCS reads were converted to HiFi reads using bamtools v2.5.1 with a minimum read quality threshold of 0.99 (Barnett et al. 2021). Adapter contamination was removed using HiFiAdapterFilt (minimum 44 bp, 97% match), which removed 1,768 reads (0.067%) and retained 2,645,236 reads (Sim et al. 2022).

To remove host plant and microbial contamination, retained reads were classified using a custom Kraken2 database (Wood et al. 2019) containing: (1) the *Coffea arabica* RefSeq genome assembly (GCF_003713225.1), (2) all available bacterial genomes from the Kraken2 standard database, (3) representative fungal genomes, and (4) published weevil genomes including *Cosmopolites sordidus,* the remainder of the genomes were the same as in, Rodriguez Ruiz & Van Dam (2023). Reads classified as Hexapoda were retained using seqkit grep and used for downstream assembly. This three-tiered metagenomic approach, combining (1) pre-assembly read classification, (2) post-assembly haplotig purging and contaminant removal, and (3) independent assembly and deep learning taxonomic classification of the filtered fraction (see Metagenomic binning below), follows the pipeline established for the banana weevil *C. sordidus* genome (Rodriguez Ruiz & Van Dam 2023), where pre-assembly Kraken2 classification of HiFi reads reduced bacterial gene contamination by 6.5-fold compared to post-assembly filtering alone (179 vs. 1,158 bacterial-annotated genes), demonstrating that removing microbial and host plant reads prior to the HiFiasm assembly phase is essential for producing clean beetle genomes from specimens with complex associated microbiomes. The same custom Kraken2 database architecture containing all available bacterial genomes, representative fungal genomes, and related Curculionoidea genomes was used here, with the addition of the *Coffea arabica* genome assembly to filter host plant contamination.

Filtered reads were assembled using HiFiasm v0.16 (Cheng et al. 2021, 2022) with 120 threads and level 3 aggressive haplotig purging (−l 3). Haplotigs and heterozygous overlaps were further removed using purge_dups (Guan et al. 2020) with default parameters. Assembly statistics were generated using bbmap stats.sh (Bushnell 2014), and completeness was assessed using BUSCO v5.3.0 in genome mode against the insecta_odb10 lineage dataset (Simão et al. 2015).

### Contamination visualization and bacterial scaffold separation

Taxonomic composition of the assembly was visualized using BlobToolKit (Laetsch and Blaxter 2017; Challis et al. 2020), generating GC-coverage blob plots and snail plots for assembly quality assessment. BLAST similarity searches against the NCBI nucleotide database were used to assign taxonomic affiliations to individual scaffolds. Scaffolds classified as non-arthropod were separated into a bacterial scaffold assembly for independent analysis.

### Metagenomic binning and taxonomic classification of bacterial scaffolds

Reads that failed the Kraken2 arthropod classification filter were independently assembled using HiFiasm to produce a bacterial scaffold assembly (16,216 contigs, 295 Mb). This assembly was analyzed using VAMB v4 (Nissen et al. 2021), a variational autoencoder-based metagenomic binner that clusters contigs by tetranucleotide frequency and coverage depth. VAMB was run with minimum bin size 50 (-b 50), producing 8,060 metagenomic bins. Taxonomic classification of individual contigs was performed using TaxVAMB (Kutuzova et al. 2026) with its built-in Taxometer module (Kutuzova et al. 2024), which assigns NCBI taxonomy to contigs using a deep learning classifier trained on reference genomes. Bin quality was assessed using CheckM2 (Chklovski et al. 2023). All twelve VAMB parameter combinations tested (minimum bin sizes 50–200) produced statistically equivalent levels of contamination (median <5%), so results are reported for b = 50 only. Coverage was computed from read mapping with minimap2 (Li 2018). This two-stage approach (assembly followed by compositional binning and deep learning taxonomy) provides comprehensive interrogation of the microbial community associated with *A. fasciculatus* specimens, and is critical for testing whether bacterial caffeine degradation genes might reside in the associated microbiome rather than in the beetle genome itself.

### Repeat masking and gene prediction

Transposable elements and repetitive sequences were identified *de novo* using RepeatModeler2 (Flynn et al. 2020) and masked using RepeatMasker v4.1.5 (Smit et al. 2015) with rmblastn v2.14.1. Gene prediction was performed on the repeat-masked assembly using BRAKER2 v2.1.6 (Brůa et al. 2021) in EP mode, with the insecta_odb10 reference protein database as extrinsic evidence. BRAKER2 employed GeneMark-EP for initial gene finding and AUGUSTUS for final gene model optimization. Predicted proteins were functionally annotated through DIAMOND BLASTp (Buchfink et al. 2015) against the SwissProt database.

### Identification of cytochrome P450 genes

Predicted proteins with SwissProt annotations matching “cytochrome P450” or “CYP” were extracted to identify the *A. fasciculatus* P450 complement. These 92 P450 proteins were searched against the three *Drosophila melanogaster* P450s experimentally demonstrated to metabolize caffeine: CYP12D1-d (AAG22287.3), CYP6d5 (AAF55009.1), and CYP6a8 (AAF58185.2) (Coelho et al. 2015), using BLASTp v2.17.0 (Camacho et al. 2009; default parameters, e-value threshold 1 × 10^-5^). Genomic organization of P450 genes was examined by mapping gene coordinates from the GFF3 annotation to identify tandem gene clusters, defined as two or more P450 genes within 200 kb on the same scaffold.

### Searches for bacterial caffeine N-demethylase genes

To test whether *A. fasciculatus* harbors bacterial-origin caffeine degradation genes, we conducted four comprehensive BLAST searches using the bacterial N-demethylase proteins from *Pseudomonas putida* CBB5 as queries: NdmA (UniProt: H9N289), NdmB (UniProt: H9N290), NdmC (UniProt: F0E1K6), and NdmD (UniProt: H9N291) (Summers et al. 2012). The searches were: (1) tBLASTn of NdmA/B/C/D against the *A. fasciculatus* genome assembly; (2) tBLASTn against the separated bacterial scaffold assembly (16,216 contigs); (3) BLASTp against the complete *A. fasciculatus* predicted proteome (22,684 proteins); and (4) BLASTp against the *Drosophila* caffeine P450 database as a control. All searches used e-value threshold 1 × 10^-5^.

### Molecular docking

To compare the caffeine-binding properties of insect P450s and bacterial N-demethylases, molecular docking was performed using AutoDock Vina v1.2.7 (Eberhardt et al. 2021). Reference protein structures were obtained from the AlphaFold Protein Structure Database v6 (Jumper et al. 2021) for *D. melanogaster* CYP12D1-p, CYP6d5, and CYP6a8, and from the Protein Data Bank (PDB: 6ICP) for the *P. putida* NdmA crystal structure with bound caffeine (Kim et al. 2019). Receptor structures were prepared using Open Babel (O’Boyle et al. 2011) and caffeine (SMILES: CN1C=NC2=C1C(=O)N(C(=O)N2C)C) was prepared as a 3D ligand using RDKit (Landrum 2024) and Meeko v0.7.1 (Santos-Martins et al. 2025). Docking was performed with exhaustiveness = 32, a 30 Å cubic search box centered on each protein, and nine binding poses.

### Protein structure prediction

Three-dimensional structures of the top *A. fasciculatus* P450 candidates were predicted using ColabFold v1.6.1 (Mirdita et al. 2022), which implements AlphaFold2 (Jumper et al. 2021) with MMseqs2 for multiple sequence alignment generation. Structure quality was assessed by predicted local distance difference test (pLDDT) and predicted template modeling (pTM) scores. Conserved P450 structural motifs were identified by regular expression matching: the heme-binding cysteine (FXXGXXXCXG), EXXR helix interaction motif, PERF motif, and I-helix acid-alcohol pair.

### Structural homology search

To independently confirm the fold classification of the top *Araecerus* P450 candidates, predicted structures were searched against all experimentally solved structures in the Protein Data Bank using Foldseek v10.941cd33 (van Kempen et al. 2024). Each ColabFold-predicted structure was queried against the PDB100 database (clustered at 100% sequence identity) using the easy-search workflow with default parameters, an E-value threshold of 1 × 10^-5^, and a maximum of 20 hits per query. This structure-based approach complements the sequence-based BLAST searches by detecting shared three-dimensional folds regardless of sequence divergence, and provides a stringent test of whether the *Araecerus* candidates adopt a cytochrome P450 fold or a bacterial Rieske-type non-heme iron oxygenase fold.

### Structural superposition

Structural similarity between ColabFold-predicted *Araecerus* P450 structures and reference insect P450s was assessed using the PyMOL align command (Schrödinger, LLC), which performs sequence-dependent structural superposition with iterative outlier rejection and reports root-mean-square deviation (RMSD) over aligned Cα atoms. Each *Araecerus* candidate was superimposed on the *Drosophila melanogaster* CYP12D1-p AlphaFold v6 structure.

### Phylogenetic analysis of P450 candidates

To determine whether the A. fasciculatus caffeine P450 candidates represent lineage-specific expansions or ancestral genes predating the seed-feeding transition, we constructed a multi-species cytochrome P450 phylogeny. The four top *Araecerus* candidates were searched against the chromosome-level genomes of two non-seed-feeding Anthribidae from the Darwin Tree of Life project (*Pseudeuparius sepicola* (GCA_963920635.1, Booth et al. 2024) and *Platystomos albinus* (GCA_964106875.1, Crowley et al. 2025)) using tBLASTn v2.17.0 (e-value threshold 1 × 10^-20^, maximum 10 targets per query). Since neither DToL genome includes gene predictions, translated subject sequences from the best two hits per query per species were extracted directly from the BLAST output. Reference P450 protein sequences were obtained from NCBI for representative insect CYP families: CYP6 (*Tribolium castaneum* CYP6BQ9, *Dendroctonus ponderosae* CYP6DE1, *Leptinotarsa decemlineata* CYP6BJ1, *Musca domestica* CYP6D3, *Bombyx mori* CYP6AE2), CYP9 (*D. melanogaster* CYP9B2, B. mori CYP9A19), CYP4 (*D. melanogaster* CYP4E2, T. castaneum CYP4Q7), and the three *D. melanogaster* caffeine-metabolizing P450s (CYP12D1, CYP6a8, CYP6d5; Coelho et al. 2015). Saccharomyces cerevisiae CYP51/ERG11 was included as an outgroup. Sequences shorter than 250 amino acids were excluded. Multiple sequence alignment was performed with MAFFT v7.526 (Katoh and Standley 2013) using the --auto strategy, followed by automated trimming with trimAl (Capella-Gutiérrez et al. 2009). Phylogenetic inference was performed with IQ-TREE v3.0.1 (Minh et al. 2020) using ModelFinder (Kalyaanamoorthy et al. 2017) for model selection, 1,000 ultrafast bootstrap replicates (Hoang et al. 2018), and 1,000 SH-aLRT replicates. The tree was rooted on CYP51.

## Results

### Sequencing Results

PacBio HiFi sequencing yielded 2,647,004 circular consensus sequence (CCS) reads totaling 21.0 Gb. Reads had a mean length of 7,920 bp (median 7,990 bp; N50 9,380 bp; range 42–36,014 bp).

### Genome assembly

We evaluated three-step genome assembly pipeline by progressively increasing stringency (Table 1). The initial unfiltered assembly from all 2,647,004 CCS reads yielded 262 contigs spanning 687 Mb with the highest BUSCO completeness (99.5%) but also the highest duplication rate (19.5%), indicating substantial retention of haplotigs and potential contaminant sequences. The second assembly, using reads filtered through HiFiAdapterFilt and Kraken2 metagenomic classification, produced 5,822 contigs spanning 626 Mb with 94.4% completeness and 19.2% duplication. The third and final assembly added purge_dups processing, reducing duplication to 3.1% at the cost of some completeness (88.5%), with 3,617 scaffolds spanning 475 Mb (Table 1). The dramatic reduction in duplication (19.5% to 3.1%) with only a moderate decrease in completeness confirmed this as the optimal assembly for downstream analysis.

**Table 1.** Comparison of three *Araecerus fasciculatus* genome assembly strategies.

| Metric | Unfiltered | Kraken2-filtered | Final (+ purge_dups) |
| --- | --- | --- | --- |
| Contigs/scaffolds | 262 | 5,822 | 3,617 |
| Total size (Mb) | 687 | 626 | 475 |
| N50 (kb) | 8,409 | 170 | 170 |
| Longest scaffold (kb) | 24,251 | 924 | 924 |
| BUSCO complete (%) | 99.5 | 94.4 | 88.5 |
| BUSCO single-copy (%) | 80.0 | 75.2 | 85.4 |
| BUSCO duplicated (%) | 19.5 | 19.2 | 3.1 |

BlobToolKit visualization of the final assembly (Fig. 1) showed a single dominant cluster of scaffolds with GC content centered at ∼35.6% and consistent read coverage, characteristic of a haploid arthropod genome. The assembly snail plot (Fig. 2) confirmed an N50 of 170 kb with the longest scaffold at 924 kb. A small number of scaffolds with divergent GC content or anomalous coverage were identified as probable bacterial contaminants and segregated into a separate bacterial scaffold assembly for independent analysis (Fig. 3).

**Figure 1.**
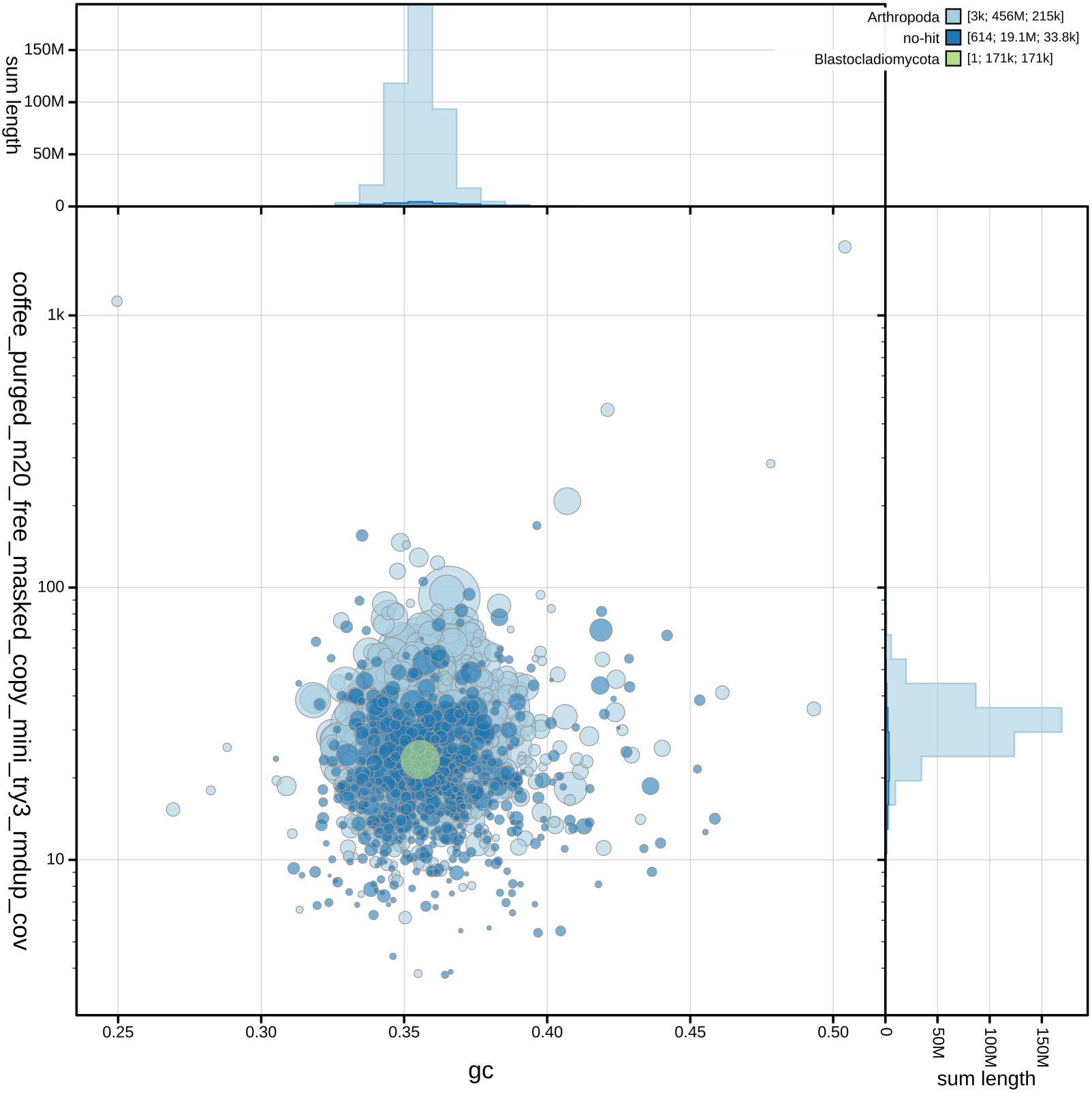
Genome assembly overview of *Araecerus fasciculatus* from Puerto Rico. BlobToolKit Taxon-annotated GC-coverage blob plot. Each circle represents a scaffold with size proportional to length. The main cluster (GC ∼35%, coverage ∼15-30x) represents the beetle genome.

**Figure 2.**
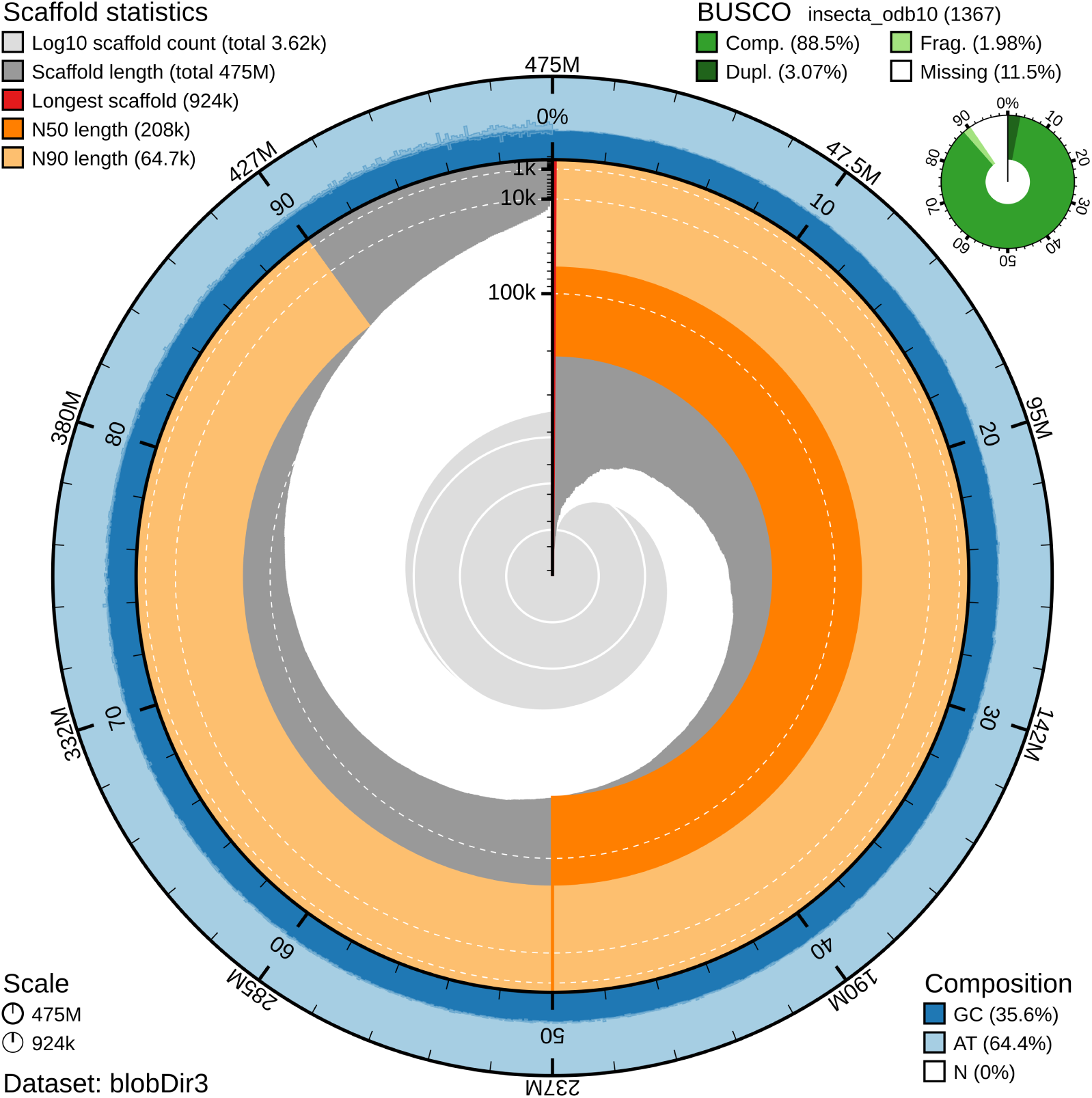
BlobToolKit snail plot of the *Araecerus* fasciculatus from Puerto Rico genome assembly showing scaffold statistics: 3,617 scaffolds totaling 475 Mb, longest scaffold 924 kb, N50 = 170 kb. Inset circle shows BUSCO completeness (green = complete 88.5%, orange = fragmented 2.0%, red = missing 9.5%).

**Figure 3.**
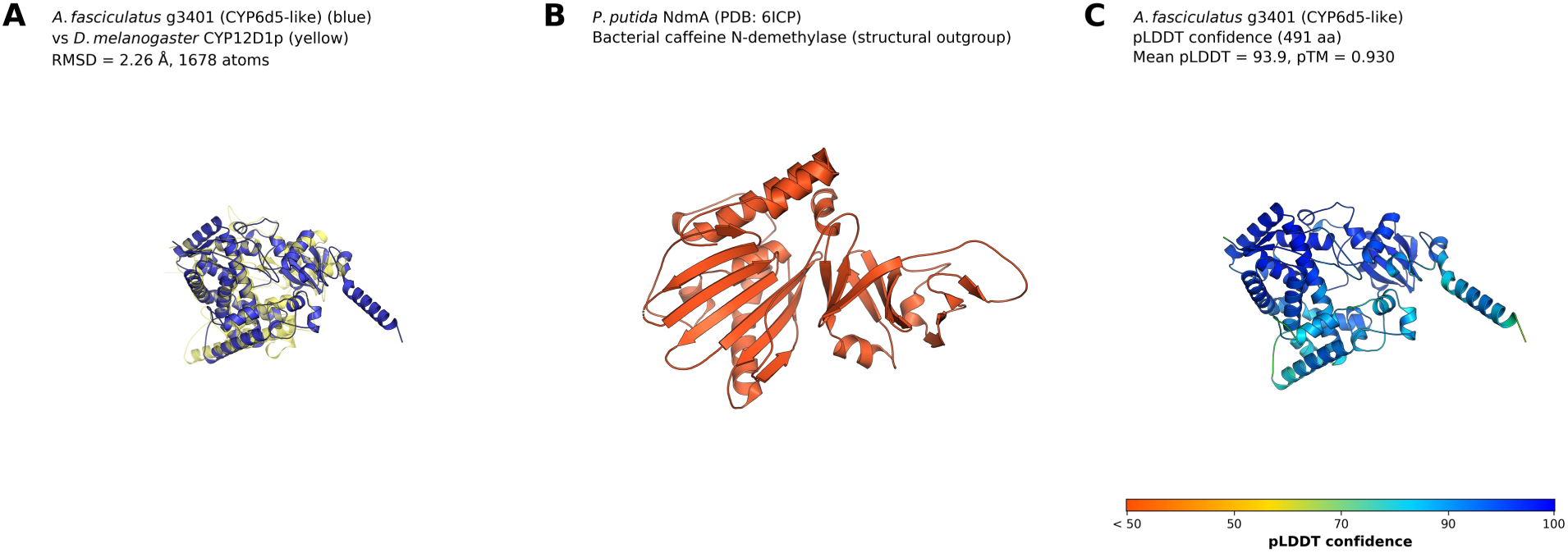
AlphaFold2 predicted structures of *Araecerus fasciculatus* P450 candidates compared to *Drosophila* CYP12D1-p and bacterial NdmA. For each candidate (g3401, g9600, g17057, g11033): (A) structural superposition of the *Araecerus* candidate (blue) with *D. melanogaster* CYP12D1-p (yellow, semi-transparent), showing conserved P450 fold architecture with RMSD values; (B) the bacterial P. putida NdmA (PDB: 6ICP, orange) as a structural outgroup, illustrating the completely different Rieske-type beta-sheet-dominated fold; (C) the *Araecerus* candidate colored by AlphaFold2 pLDDT confidence (blue > 90, very high; cyan 70–90, confident; yellow 50–70, low; orange < 50, very low). Best candidate (g3401, pLDDT = 93.9) shown as main figure; remaining candidates (g9600, g17057, g11033) as supplementary figures.

### Metagenomic analysis of associated bacterial community

Independent assembly of reads that failed arthropod classification produced 16,216 contigs spanning 295 Mb. VAMB metagenomic binning generated 8,060 bins, and TaxVAMB Taxometer classified contigs into four major categories (Table S3): Bacteria (8,069 contigs, 49.8%), Eukaryota (4,464 contigs, 27.5%), unresolved “cellular organisms” (2,243 contigs, 13.8%), and unclassified (1,440 contigs, 8.9%).

The eukaryotic fraction was dominated by weevil-origin sequences: *Sitophilus oryzae* (1,003 contigs), *Listronotus bonariensis* (1,527), *Pachyrhynchus sulphureomaculatus* (792), and *Rhynchophorus ferrugineus* (222)—all Curculionidae. These represent *A. fasciculatus* genomic DNA whose closest database matches are other weevil genomes, confirming the phylogenetic isolation of Anthribidae in current reference databases. An additional 729 contigs were classified as *Coffea arabica* (host plant contamination) and 42 as *Musa acuminata*.

The bacterial fraction contained a diverse assemblage of common environmental and gut-associated bacteria. The most abundant bacterial taxa were *Romboutsia hominis* (Peptostreptococcaceae, 208 contigs), *Paenibacillus swuensis* (Paenibacillaceae, 174), *Campylobacter lari* (Campylobacteraceae, 170), *Companilactobacillus paralimentarius* (Lactobacillaceae, 104), *Clostridium perfringens* (Clostridiaceae, 79), and *Streptococcus uberis* (Streptococcaceae, 71). Several known insect endosymbionts were detected: *Wolbachia pipientis* (5 contigs, 110 kb total), *Sodalis* endosymbiont (2 contigs, 59 kb), and *Wigglesworthia glossinidia* (2 contigs, 33 kb), the latter two likely representing database misassignments of a true *Nardonella*-like Curculionoidea endosymbiont not yet represented in reference databases. The presence of *Wolbachia* is consistent with reports of this reproductive parasite in other Curculionoidea.

Critically, although 34 contigs were classified as *Pseudomonas* spp. within the Pseudomonadota (1,833 total Pseudomonadota contigs), the tBLASTn search for caffeine N-demethylase genes (ndmA/B/C/D from *P. putida* CBB5) against this entire 16,216-contig bacterial assembly returned zero hits, confirming the complete absence of bacterial caffeine degradation genes from the *A. fasciculatus*-associated microbiome (Fig. 4).

**Figure 4.**
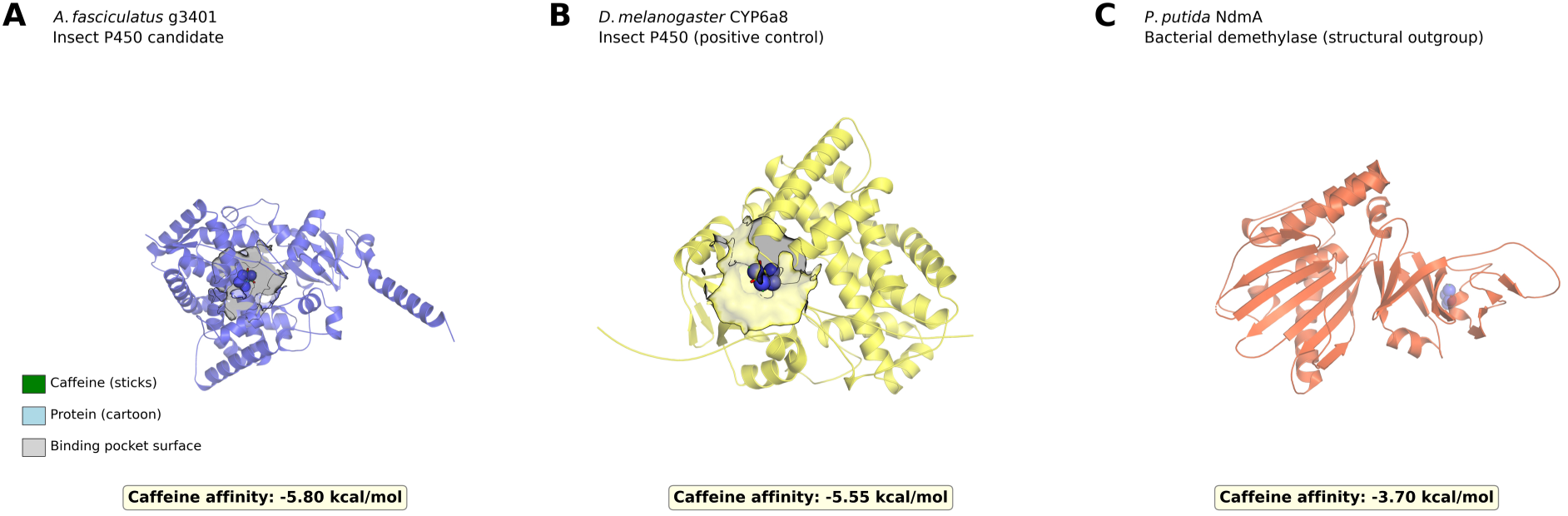
Molecular docking of caffeine to *A. fasciculatus* P450 candidates, *Drosophila* CYP6a8 (positive control), and bacterial NdmA (structural outgroup). For each candidate: (A) *Araecerus* P450 (blue cartoon, semi-transparent) with docked caffeine (green sticks) and binding pocket surface (transparent); (B) *D. melanogaster* CYP6a8 (yellow) with caffeine; (C) P. putida NdmA (orange) with caffeine. Binding affinities (kcal/mol) labeled below each panel. Best candidate g3401 (−5.80 kcal/mol) shown as main figure; g9600 (−5.63), g17057 (−5.55), g11033 (−5.41) as supplementary.

### Repeat content and gene prediction

RepeatMasker identified repetitive elements occupying 306.2 Mb (64.47%) of the 475 Mb assembly (Table S1). Post hoc classification of the 4,095 de novo repeat consensus sequences by BLASTx against the Dfam RepeatPeps protein database (e-value < 1e-5) identified protein-coding domains in 557 families (13.6%), which accounted for 121,440 genomic elements spanning 59.3 Mb (12.5% of the assembly). LTR retrotransposons were the most abundant classified category (26.1 Mb, 5.5%), followed by LINEs (17.6 Mb, 3.7%) and DNA transposons (13.3 Mb, 2.8%). The remaining 3,538 families (297.4 Mb, 62.6%) lacked recognizable protein domains, consistent with non-autonomous elements, highly degraded copies, or lineage-specific repeats absent from curated databases. The overall repeat landscape of this anthribid genome is largely composed of novel, lineage-specific transposable elements.

BRAKER2 gene prediction on the repeat-masked assembly identified 22,384 protein-coding genes producing 22,684 transcripts. BUSCO assessment of the predicted protein set in protein mode (insecta_odb10) yielded 85.5% complete (78.9% single-copy, 6.6% duplicated), 3.4% fragmented, and 11.1% missing (Table S2), consistent with the genome-level BUSCO and indicating that the majority of conserved insect genes were successfully predicted.

Functional annotation via SwissProt BLASTp assigned putative functions to 11,783 transcripts (51.9%), of which 1,368 (11.6%) were transposable element-derived proteins, consistent with the high repeat content. The remaining 10,415 functionally annotated transcripts represented putative functional protein coding genes, while 10,901 transcripts were classified as proteins of unknown function (Table 2).

**Table 2.** Functional annotation summary of 22,684 predicted transcripts.

| Category | Count | % of total |
| --- | --- | --- |
| Total predicted transcripts | 22,684 | 100.0 |
| SwissProt annotated | 11,783 | 51.9 |
| TE-derived proteins | 1,368 | 6.0 |
| Functional proteins | 10,415 | 45.9 |
| Unknown function | 10,901 | 48.1 |
| Detoxification enzymes | 324 | — |
| Esterases/carboxylesterases | 105 | — |
| Cytochrome P450s | 94 | — |
| UDP-glycosyltransferases | 56 | — |
| ABC transporters | 36 | — |
| Glutathione S-transferases | 33 | — |
| Digestive enzymes | 353 | — |
| Developmental genes | 167 | — |
| Chemosensory receptors | 39 | — |

The annotated gene set included a substantial detoxification gene repertoire totaling 324 transcripts across five major enzyme families: esterases/carboxylesterases (105), cytochrome P450s (94), UDP-glycosyltransferases (56), ABC transporters (36), and glutathione S-transferases (33). Additional functional categories included digestive enzymes (353 transcripts: trypsins, cathepsins, proteases, lipases, amylases), developmental genes (167: juvenile hormone esterase, ecdysteroid-related, cuticle proteins, chitin-binding), and chemosensory receptors (39: odorant, gustatory, and ionotropic receptors) (Table 2). The large detoxification gene complement is consistent with the polyphagous feeding ecology of *A. fasciculatus* across >100 chemically diverse stored products.

### Expanded cytochrome P450 repertoire

SwissProt annotation identified 92 cytochrome P450 genes in the *A. fasciculatus* genome, representing approximately 0.41% of the predicted gene complement. This count is comparable to other Coleoptera: *T. castaneum* has 143 CYP genes, *D. ponderosae* has 86, and the average in Coleoptera is approximately 100–150. BLASTp comparison of all 92 *Araecerus* P450 proteins against the three *Drosophila* caffeine-metabolizing P450s yielded 292 hits at e-value < 1 × 10^-5^ (Table 3). The top six candidates showed strong similarity to one or more *Drosophila* caffeine P450s (see Table 3).

**Table 3.** Top six *Araecerus fasciculatus* P450 candidates from BLASTp comparison against Drosophila caffeine-metabolizing P450s.

| Gene | Scaffold | Annotation | Best Match | Identity | E-value |
| --- | --- | --- | --- | --- | --- |
| g15909 | ptg003214l | CYP12a5-like | Cyp12d1-d | 37.4% | 9.7e-110 |
| g11033 | ptg000464l | CYP6a13-like | CYP6a8 | 38.6% | 3.4e-105 |
| g17057 | ptg000257l | CYP49a1-like | Cyp12d1-d | 33.9% | 1.0e-104 |
| g9600 | ptg001747l | CYP6K1-like | CYP6a8 | 34.9% | 2.7e-101 |
| g3401 | ptg001867l | CYP6K1-like | CYP6d5 | 33.7% | 5.4e-101 |
| g11030 | ptg000464l | CYP6a13-like | CYP6a8 | 37.7% | 1.7e-100 |

Two notable tandem P450 gene clusters were identified: **Cluster 1** (scaffold ptg000464l): four CYP6a13-like genes (g11030, g11033, g11034, g11035) spanning a 71 kb region (15– 86 kb), all annotated as similar to *Drosophila* Cyp6a13. **Cluster 2** (scaffold ptg001867l): five CYP6K1-like genes (g3400, g3401, g3402, g3403, g3404) spanning a 49 kb region (141– 190 kb). Tandem gene clusters are a hallmark of adaptive P450 gene family expansion in insects, commonly associated with xenobiotic detoxification (Feyereisen 2012). The presence of two such clusters in *A. fasciculatus*, both containing genes with strong homology to *Drosophila* caffeine-metabolizing P450s, suggests lineage-specific expansion of detoxification capacity.

### Absence of bacterial caffeine N-demethylase genes

All four BLAST searches for bacterial caffeine N-demethylase genes returned zero hits (Table 4). The *Pseudomonas putida* CBB5 NdmA, NdmB, NdmC, and NdmD proteins produced no matches at any e-value threshold against: (1) the *A. fasciculatus* genome (tBLASTn), (2) the separated bacterial scaffold assembly of 16,216 contigs (tBLASTn), (3) all 22,684 predicted *A. fasciculatus* proteins (BLASTp), or (4) the extracted *Araecerus* P450 proteins (BLASTp, as expected for a negative control). This comprehensive negative result demonstrates that neither the beetle genome nor its associated bacterial community encodes recognizable caffeine N-demethylase homologs, in stark contrast to the Scolytinae system where such genes are present both in the beetle genome (as HGT) and in associated bacteria.

**Table 4.** Comprehensive BLAST searches for bacterial caffeine N-demethylase (ndm) genes.

| Search | Query | Database | Method | Hits |
| --- | --- | --- | --- | --- |
| 1 | NdmA/B/C/D | *A. fasciculatus* genome | tBLASTn | 0 |
| 2 | NdmA/B/C/D | Bacterial scaffolds (16,216 contigs) | tBLASTn | 0 |
| 3 | NdmA/B/C/D | *A. fasciculatus* proteome (22,684) | BLASTp | 0 |
| 4 | NdmA/B/C/D | *Drosophila* caffeine P450s (control) | BLASTp | 0 |

### Molecular docking

Molecular docking of caffeine against ColabFold-predicted *Araecerus* P450 structures and reference proteins revealed that all four beetle candidates exhibit predicted caffeine-binding affinities comparable to or exceeding experimentally validated *Drosophila* caffeine-metabolizing P450s (Table 5). The strongest predicted binding was for *Araecerus* g3401 (−5.80 kcal/mol), followed by g9600 (−5.63 kcal/mol), g17057 (−5.55 kcal/mol), and g11033 (−5.41 kcal/mol). For comparison, the *Drosophila* CYP6a8 positive control yielded −5.55 kcal/mol and CYP12D1-p yielded −5.43 kcal/mol. Three of four *Araecerus* candidates thus showed equal or stronger predicted caffeine binding than the experimentally validated *Drosophila* P450s.

**Table 5.** AutoDock Vina molecular docking results for caffeine binding to *Araecerus* P450 candidates and reference structures.

| Protein | Species | Annotation | Structure source | Affinity (kcal/mol) |
| --- | --- | --- | --- | --- |
| g3401 | *A. fasciculatus* | CYP6K1-like | ColabFold | -5.80 |
| g9600 | *A. fasciculatus* | CYP6K1-like | ColabFold | -5.63 |
| g17057 | *A. fasciculatus* | CYP49a1-like | ColabFold | -5.55 |
| g11033 | *A. fasciculatus* | CYP6a13-like | ColabFold | -5.41 |
| CYP6a8 | *D. melanogaster* | — | AlphaFold DB v6 | -5.55 |
| CYP12D1-p | *D. melanogaster* | — | AlphaFold DB v6 | -5.43 |
| CYP6d5 | *D. melanogaster* | — | AlphaFold DB v6 | -3.83 |
| NdmA (6ICP) | *P. putida* CBB5 | — | PDB crystal | -3.70 |

The bacterial N-demethylase NdmA crystal structure (PDB: 6ICP), included as a structural outgroup representing the fundamentally different Rieske-fold caffeine degradation pathway, yielded the weakest predicted binding (−3.70 kcal/mol) despite being a known caffeine-degrading enzyme. This difference reflects the distinct binding mechanisms of heme-containing P450 monooxygenases versus Rieske-type non-heme iron oxygenases: the bacterial enzymes employ a different catalytic architecture (mononuclear non-heme iron center with a [2Fe-2S] Rieske cluster) that is not well captured by the AutoDock Vina scoring function optimized for conventional binding pockets. The NdmA result should not be interpreted as indicating weaker caffeine metabolism by this enzyme, but rather illustrates the fundamental structural and mechanistic divergence between the two caffeine degradation pathways.

### Protein structure prediction of Araecerus P450 candidates

AlphaFold2 structure prediction via ColabFold v1.6.1 (3 models, 3 recycles per candidate) produced high-confidence structures for all four top *A. fasciculatus* P450 candidates (Table S4). The best-ranked model for each candidate was model 3 in all cases. Candidate g3401 (491 aa, CYP6K1-like, from the five-gene tandem cluster on ptg001867l) achieved the highest confidence (pLDDT = 93.9, pTM = 0.930), followed by g9600 (497 aa, CYP6K1-like; pLDDT = 92.8, pTM = 0.918), g17057 (518 aa, CYP49a1-like; pLDDT = 87.4, pTM = 0.879), and g11033 (613 aa, CYP6a13-like; pLDDT = 84.5, pTM = 0.735). The lower pTM score for g11033 is consistent with its unusually long sequence (613 aa vs. ∼490–520 aa for canonical P450s), suggesting a possible tandem P450 domain or N-terminal extension.

Structural superposition of the predicted *Araecerus* structures against the *Drosophila* CYP12D1-p AlphaFold reference confirmed that all four candidates adopt the canonical cytochrome P450 fold (Table S4). The closest structural match was g17057 (RMSD = 1.51 Å over 2,637 aligned atoms), indicating near-identical backbone topology. Candidates g9600 (RMSD = 2.24 Å, 1,513 atoms) and g3401 (RMSD = 2.26 Å, 1,678 atoms) also showed excellent structural correspondence. Candidate g11033 showed somewhat greater divergence (RMSD = 3.48 Å, 2,070 atoms), with the additional aligned atoms reflecting its extended structure. For comparison, the bacterial NdmA (PDB: 6ICP) adopts a completely different Rieske-type non-heme iron oxygenase fold dominated by β-sheets, which is structurally unrelated to the all-α-helical P450 architecture.

Foldseek structural homology searches against all experimentally solved structures in the Protein Data Bank (PDB100) provided independent, structure-based confirmation that the *Araecerus* candidates are genuine cytochrome P450s (Table S5). All 80 hits across the four candidates (20 per candidate) were cytochrome P450 enzymes, with zero hits to any bacterial caffeine N-demethylase, Rieske-fold oxygenase, or any other non-P450 enzyme fold. All hits achieved probability scores of 1.000, confirming unambiguous fold classification. The top structural matches were predominantly human drug-metabolizing P450s: CYP3A4 (PDB: 5TE8, TM-score 0.91 for g3401; TM-score 0.90 for g9600), CYP3A7 (PDB: 8GK3, TM-score 0.90 for g11033), and CYP11A1 (PDB: 3MZS, TM-score 0.92 for g17057). These results confirm by three-dimensional structure, independent of sequence similarity, that the *Araecerus* caffeine detoxification candidates are bona fide cytochrome P450 enzymes structurally unrelated to the bacterial ndm caffeine degradation pathway.

P450 conserved motif analysis confirmed functional hallmarks in the top candidates. Candidate g3401 possesses all four diagnostic P450 motifs: the heme-binding cysteine (FXXGXXXCXG), the EXXR helix interaction motif, the PERF motif, and the I-helix acid-alcohol pair. Candidate g9600 likewise retains all four motifs. Candidate g17057 retains three motifs (heme Cys, EXXR, PERF), while g11033 retains three (heme Cys, EXXR, I-helix acid-alcohol pair). The retention of the critical heme-binding cysteine in all four candidates is consistent with catalytically competent P450 enzymes.

### Phylogenetic placement of Araecerus P450 candidates

The P450 phylogeny (30 sequences, 358 trimmed alignment positions, LG+I+G4 model) revealed that all four *A. fasciculatus* caffeine P450 candidates have clear orthologs in both non-seed-feeding DToL anthribids (Fig. 5). tBLASTn searches recovered strong hits in *Pseudeuparius sepicola* for all four candidates (g3401: 61.8% identity, e = 5.25 × 10^-178^; g11033: 39.6%, e = 1.61 × 10^-105^; g9600: 35.9%, e = 1.20 × 10^-94^; g17057: 35.6%, e = 1.35 × 10^-41^), and similarly strong hits in *Platystomos albinus* (g3401: 69.1%; g9600: 68.4%; g11033: 42.2%; g17057: 36.0%). These orthologs placed sister to their respective *Araecerus* candidates with strong support in the phylogeny, confirming orthology rather than paralogy.

**Figure 5.**
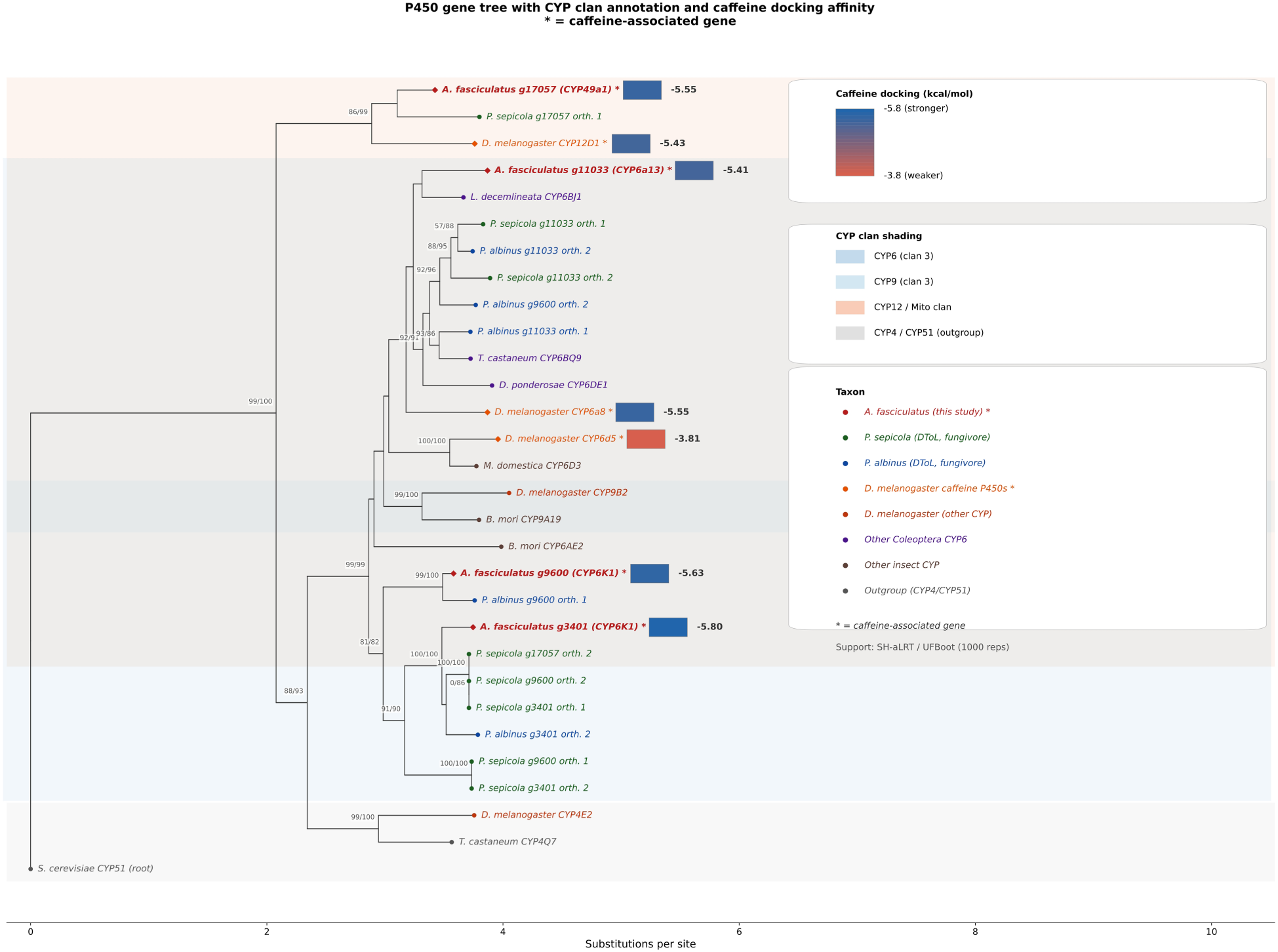
Cytochrome P450 phylogeny of *Araecerus* fasciculatus caffeine detoxification candidates. Maximum likelihood tree (IQ-TREE 3, LG+I+G4 model) of 30 P450 protein sequences including the four *Araecerus* candidates (red), orthologs from two non-seed-feeding DToL Anthribidae (Pseudeuparius sepicola, green; Platystomos albinus, blue), *Drosophila* caffeine-metabolizing P450s (orange), representative beetle CYP6 enzymes (purple), and CYP4/CYP9/CYP51 family representatives. Node support values are SH-aLRT / ultrafast bootstrap (1,000 replicates each). The tree is rooted on S. cerevisiae CYP51 (ERG11). All four *Araecerus* candidates have orthologs in both non-seed-feeding anthribids, indicating these genes predate the seed-feeding transition.

The tree resolved three major findings. First, *Araecerus* candidates g3401 and g9600 (both CYP6K1-like) formed a well-supported anthribid-specific CYP6 subclade (99.4/99 SH-aLRT/UFBoot), with DToL anthribid orthologs nested within. Second, candidate g11033 (CYP6a13-like) placed within the broader beetle CYP6 radiation alongside *T.* castaneum CYP6BQ9 and *D.* ponderosae CYP6DE1, with the *Drosophila* caffeine P450 CYP6a8 as a more distant sister (97.6/75). Third, candidate g17057 (CYP49a1-like) placed on a long branch sister to *D. melanogaster* CYP12D1 (86.6/99), suggesting membership in the CYP12/mitochondrial P450 clan rather than the CYP6 clade occupied by the other three candidates. The CYP4 representatives formed a well-supported outgroup (99.9/100) and CYP51 rooted the tree (99.8/100).

The presence of clear orthologs in both non-seed-feeding anthribids demonstrates that these P450 genes predate the transition from fungivory to seed-feeding in *Araecerus*. The caffeine detoxification capacity thus appears to have evolved through neofunctionalization or regulatory changes in ancestral anthribid P450s, rather than through de novo gene family expansion. Furthermore, the *Araecerus* candidates predominantly belong to the CYP6 family (clan 3), while the *Drosophila* caffeine P450 CYP12D1 belongs to the mitochondrial P450 clan, confirming convergent recruitment of different P450 subfamilies for caffeine metabolism in these phylogenetically distant insect lineages.

## Discussion

The high-quality genome of *A. fasciculatus* revealed a convergent system for caffeine detoxification within insects for the first time. This study presents the first functionally annotated genome for a member of the Anthribidae, providing a critical resource for one of the most species-rich but under-represented in terms of genomic resources for families within Curculionoidea. Two chromosome-level anthribid assemblies have recently been released through the Darwin Tree of Life project (*Pseudeuparius sepicola*, 769 Mb, scaffold N50 = 68.5 Mb, BUSCO 98.5%, Booth et al. 2024; *Platystomos albinus*, 555 Mb, scaffold N50 = 69.4 Mb, BUSCO 98.8%, Crowley et al. 2025), though neither includes gene prediction or functional annotation. Our scaffold-level assembly (475 Mb, N50 = 170 kb, BUSCO 88.5%) is more fragmented than these Hi-C-scaffolded genomes, but provides the gene models and functional characterization necessary for investigating the molecular basis of caffeine detoxification. The dramatic reduction in duplication from 19.5% to 3.1% through our three-tiered metagenomic approach (pre-assembly read classification, post-assembly haplotig purging, and independent characterization of the filtered fraction via VAMB, TaxVAMB and Taxometer;(Nissen et al. 2021; Rodriguez Ruiz & Van Dam 2023; Kutuzova et al. 2024; Kutuzova et al. 2026) demonstrates the importance of stringent contamination removal for organisms with complex microbiomes.

The high repeat content (64.47%) is noteworthy. Post hoc protein-based classification revealed that LTR retrotransposons (5.5%), LINEs (3.7%), and DNA transposons (2.8%) comprise the identifiable fraction, although 62.6% of masked sequence lacked recognizable protein domains. This high proportion of unclassifiable repeats likely reflects the phylogenetic isolation of Anthribidae, with lineage-specific transposable element expansions poorly represented in current databases. Future reclassification using updated Coleoptera-specific repeat libraries may improve annotation of these elements.

### Convergent caffeine detoxification in Curculionoidea

Our results provide strong computational evidence that *A. fasciculatus* employs a fundamentally different caffeine detoxification strategy from the well-studied scolytine system. While *H. hampei* relies on horizontally transferred bacterial N-demethylase genes (Vega et al. 2015) and caffeine-degrading gut bacteria (Vega et al. 2021), *A. fasciculatus* appears to lack any recognizable ndm gene homologs. Critically, this absence was confirmed through four independent BLAST searches covering the beetle genome, predicted proteome, and the entire associated bacterial community, the latter comprehensively characterized through independent assembly (16,216 contigs, 295 Mb), VAMB metagenomic binning (8,060 bins), and TaxVAMB deep learning taxonomy. Despite identifying diverse bacteria including *Pseudomonas* species (34 contigs) in the associated microbiome, no caffeine N-demethylase homologs were detected. Instead, the beetle possesses an expanded repertoire of 92 cytochrome P450 genes, several of which show strong homology to the *Drosophila* CYP12D1, CYP6d5, and CYP6a8 genes experimentally demonstrated to metabolize caffeine (Coelho et al. 2015).

This represents a striking case of convergent evolution at multiple levels. Two beetle lineages within the same superfamily (Curculionoidea) independently evolved the ability to tolerate caffeine through completely different molecular mechanisms: horizontally transferred bacterial N-demethylase genes in Scolytinae (Vega et al. 2015) versus endogenous cytochrome P450 monooxygenases in Anthribidae (this study). Moreover, within the P450-mediated strategy, *Araecerus* and *Drosophila* appear to have convergently recruited different P450 subfamilies: our phylogeny places three of four *Araecerus* candidates (g3401, g9600, g11033) within the CYP6 family (clan 3), while the primary *Drosophila* caffeine P450 CYP12D1 belongs to the mitochondrial P450 clan. Candidate g17057 (CYP49a1-like) may share deeper ancestry with the CYP12 clan (86.6% SH-aLRT / 99% UFBoot support), suggesting that the ancestral P450 complement included members from multiple clans with potential for caffeine metabolism. Critically, all four *Araecerus* candidates have clear orthologs in two non-seed-feeding anthribids (*P. sepicola* and *P. albinus*), demonstrating that these genes predate the dietary shift to caffeine-containing seeds. The caffeine detoxification phenotype in *Araecerus* thus represents neofunctionalization or regulatory upregulation of ancestral broad-spectrum detoxification enzymes, rather than de novo gene acquisition, a fundamentally different evolutionary path from both the bacterial HGT in Scolytinae and potentially from the CYP12D1-dependent mechanism in *Drosophila*.

### Tandem P450 gene clusters as evidence of adaptive expansion

The identification of two tandem P450 gene clusters (4 CYP6a13-like genes on ptg000464l, 5 CYP6K1-like genes on ptg001867l) is particularly significant. Tandem gene duplication followed by neofunctionalization is a well-established mechanism for the evolution of insecticide resistance and xenobiotic detoxification in insects (Bass and Field 2011). The clustering of these genes, combined with their strong homology to known caffeine-metabolizing P450s, makes them prime candidates for functional validation. The CYP6 family, to which most of the tandem cluster genes belong, is among the most commonly implicated in xenobiotic metabolism across insects.

Molecular docking of caffeine against the four *Araecerus* ColabFold-predicted structures produced binding affinities of −5.41 to −5.80 kcal/mol, with three of four candidates equaling or exceeding the −5.55 kcal/mol obtained for the experimentally validated *Drosophila* CYP6a8. The best candidate, g3401, showed the strongest predicted caffeine binding of any structure tested (−5.80 kcal/mol) and possesses all four conserved P450 catalytic motifs (heme-binding cysteine, EXXR, PERF, and I-helix acid-alcohol pair). Combined with the Foldseek confirmation that all four candidates adopt cytochrome P450 folds structurally unrelated to the bacterial NdmA Rieske oxygenase, these computational results identify g3401, g9600, and g17057 as strong candidates for experimental validation of caffeine-metabolizing activity, with g3401 as the highest-priority target.

### Implications for evolutionary biology of P450 genes in Anthribidae

From an evolutionary perspective, this genome provides a molecular foundation for understanding the transition from fungivory (ancestral in Anthribidae) to seed feeding in *Araecerus* and related genera. The expansion of detoxification gene families may be a key adaptation enabling anthribid beetles to exploit chemically defended plant seeds, paralleling similar expansions documented in other seed-feeding beetle lineages such as Bruchinae. *Araecerus fasciculatus* usually oviposits on moldy stored products, and while the larvae do in fact burrow and feed on coffee seeds. It may be that adaptations for detoxifying fungi served as precursors for further use as seed feeders in the Anthribidae.

### Future directions

Functional validation of the candidate P450s identified here (g3401, g9600, g17057, g11033) should be pursued through heterologous expression in insect cell lines followed by caffeine metabolism assays to confirm demethylation activity. RNAi or CRISPR-mediated knockdown of these genes in live beetles, coupled with caffeine-supplemented diet bioassays, would establish whether they are necessary for survival on coffee substrates. Comparative transcriptomic profiling of beetles reared on caffeinated versus decaffeinated diets would further reveal whether these P450s are constitutively expressed or induced by caffeine exposure.

## Data Availability

The *Araecerus fasciculatus* genome assembly, raw HiFi reads, predicted proteins, gene annotations, BLAST search results, molecular docking data, and BlobToolKit visualizations are available from Zenodo (DOI: 10.5281/zenodo.20639466). The PacBio HiFi genome assembly have been deposited at NCBI under BioProject PRJNA1031120 (GenBank assembly accession GCA_050578095.1; BioSample SAMN37925741). Custom scripts for the docking pipeline are included in the Zenodo archive.

## Acknowledgments

We thank the staff of Café Mis Abuelos coffee farm in Mayagüez, Puerto Rico for providing access to coffee beans. Sequencing was performed at the University of Maryland Genomic Resource Center and thank Dr. Luke J. Tallon, Dr. Xuechu Ellie Zhao, and Dr. Lisa DeShong Sadzewicz (Maryland Genomics at the Institute for Genome Sciences) for PacBio library and sequencing work as well as on high molecular weight DNA extraction advice. Computational analyses were conducted on the Bridges2 system at the Pittsburgh Supercomputing Center (allocation BIO220058P). We thank the UPRM Invertebrate Collection Laboratory members for assistance with specimen curation. This work was supported by the University of Puerto Rico, Mayagüez Campus and Museum für Naturkunde Berlin.

## Author Contributions

**Alex R. Van Dam:** Conceptualization, Methodology, Software, Formal analysis, Investigation, Data curation, Writing – original draft, Writing – review & editing, Visualization, Project administration, Supervision.

**Laura V. Martinez Aponte:** Investigation, Methodology, Formal analysis, Data curation, Writing – original draft, Writing – review & editing.

**Alfredo Rodriguez Ruiz:** Investigation, Methodology, Writing – review & editing.

**Sean A. Locke:** Supervision, Writing – review & editing.

**Timothy J. Colston:** Supervision, Writing – review & editing.

## Supplementary Information

### Supplementary Tables

**Table S1.**
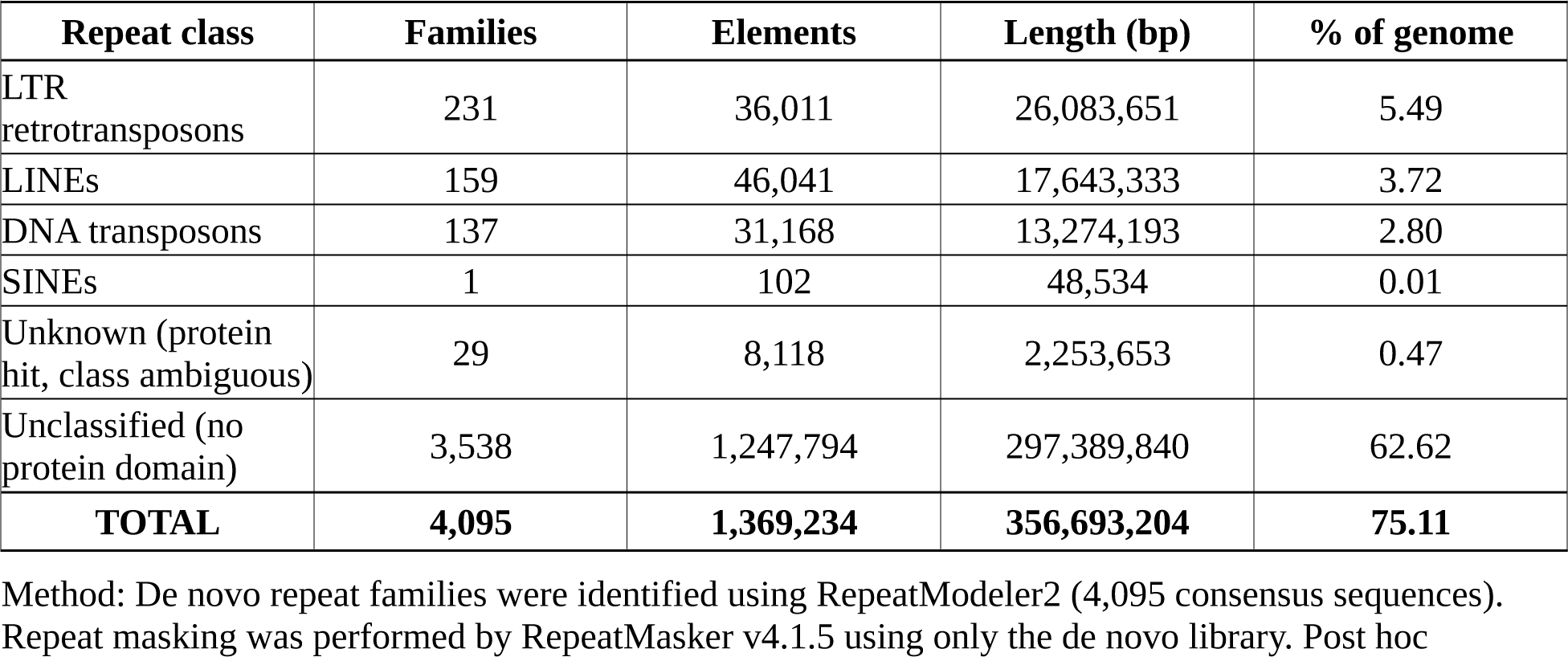
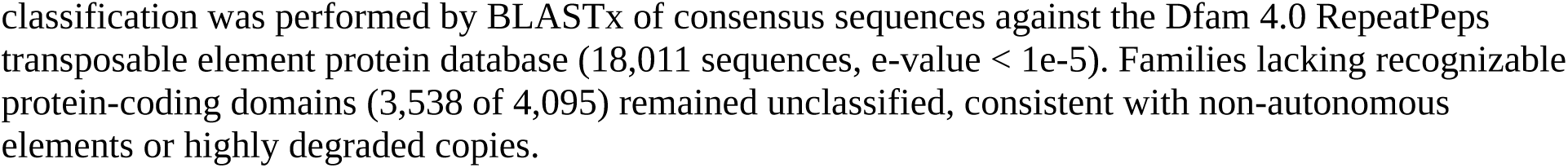
RepeatMasker summary and post hoc repeat classification for the *Araecerus fasciculatus* genome assembly.

| Repeat class | Families | Elements | Length (bp) | % of genome |
| --- | --- | --- | --- | --- |
| LTR retrotransposons | 231 | 36,011 | 26,083,651 | 5.49 |
| LINEs | 159 | 46,041 | 17,643,333 | 3.72 |
| DNA transposons | 137 | 31,168 | 13,274,193 | 2.80 |
| SINEs | 1 | 102 | 48,534 | 0.01 |
| Unknown (protein hit, class ambiguous) | 29 | 8,118 | 2,253,653 | 0.47 |
| Unclassified (no protein domain) | 3,538 | 1,247,794 | 297,389,840 | 62.62 |
| **TOTAL** | **4,095** | **1,369,234** | **356,693,204** | **75.11** |

Method: De novo repeat families were identified using RepeatModeler2 (4,095 consensus sequences). Repeat masking was performed by RepeatMasker v4.1.5 using only the de novo library. Post hoc
classification was performed by BLASTx of consensus sequences against the Dfam 4.0 RepeatPeps transposable element protein database (18,011 sequences, e-value < 1e-5). Families lacking recognizable protein-coding domains (3,538 of 4,095) remained unclassified, consistent with non-autonomous elements or highly degraded copies.

**Table S2.** *Araecerus fasciculatus* BUSCO assessment of BRAKER2 predicted proteins (insecta_odb10, protein mode). BUSCO protein-mode completeness assessment of predicted gene models.

| BUSCO category | Count | Percentage (%) |
| --- | --- | --- |
| Complete BUSCOs | 1,258 | 85.5 |
| Complete and single-copy | 1,161 | 78.9 |
| Complete and duplicated | 97 | 6.6 |
| Fragmented BUSCOs | 50 | 3.4 |
| Missing BUSCOs | 163 | 11.1 |
| Total BUSCO groups | 1,367 |  |
| Dataset | insecta_odb10 |  |
| Mode | protein |  |

**Table S3.** TaxVAMB Taxometer classification of 16,216 bacterial scaffold contigs showing domain-level composition and top species-level assignments. (attached as a csv and can also be found at https://zenodo.org/records/20639466)

**Table S4.** AlphaFold2 structure prediction and structural alignment of top A. fasciculatus P450 candidates.

| Gene | Annotation | Length (aa) | Mean pLDDT | pTM | Best model | Recycles |
| --- | --- | --- | --- | --- | --- | --- |
| g3401 | CYP6K1-like | 491 | 93.9 | 0.930 | model_3 | 3 |
| g9600 | CYP6K1-like | 507 | 92.8 | 0.920 | model_3 | 3 |
| g17057 | CYP49a1-like | 521 | 87.4 | 0.880 | model_3 | 3 |
| g11033 | CYP6a13-like | 512 | 84.5 | 0.740 | model_3 | 3 |

**The remainder of the Supplementary Tables can be found at: attached as a csv and can also be found at** https://zenodo.org/records/20639466)

**Table S5.** Foldseek structural homology search results for *A. fasciculatus* P450 candidates against PDB100.

**Table S6.** DNA extraction yields for all 12 *A. fasciculatus* specimens.

**Table S7.** Complete BLASTp results for all 92 *Araecerus* P450 proteins versus *Drosophila* caffeine P450s (292 hits).

**Table S8.** Genomic coordinates and annotations for all 92 *Araecerus* cytochrome P450 genes.

**Table S9.** P450 conserved motif analysis (heme Cys, EXXR, PERF, I-helix) for all candidate genes.

**Table S10.** Full docking results including all nine binding poses per receptor.

Tables S3 and S7 contain large datasets (16,217 and 292 rows, respectively) and are available from the Zenodo archive (DOI: 10.5281/zenodo.20639466).

### Supplementary Figures

**Figure S1.**
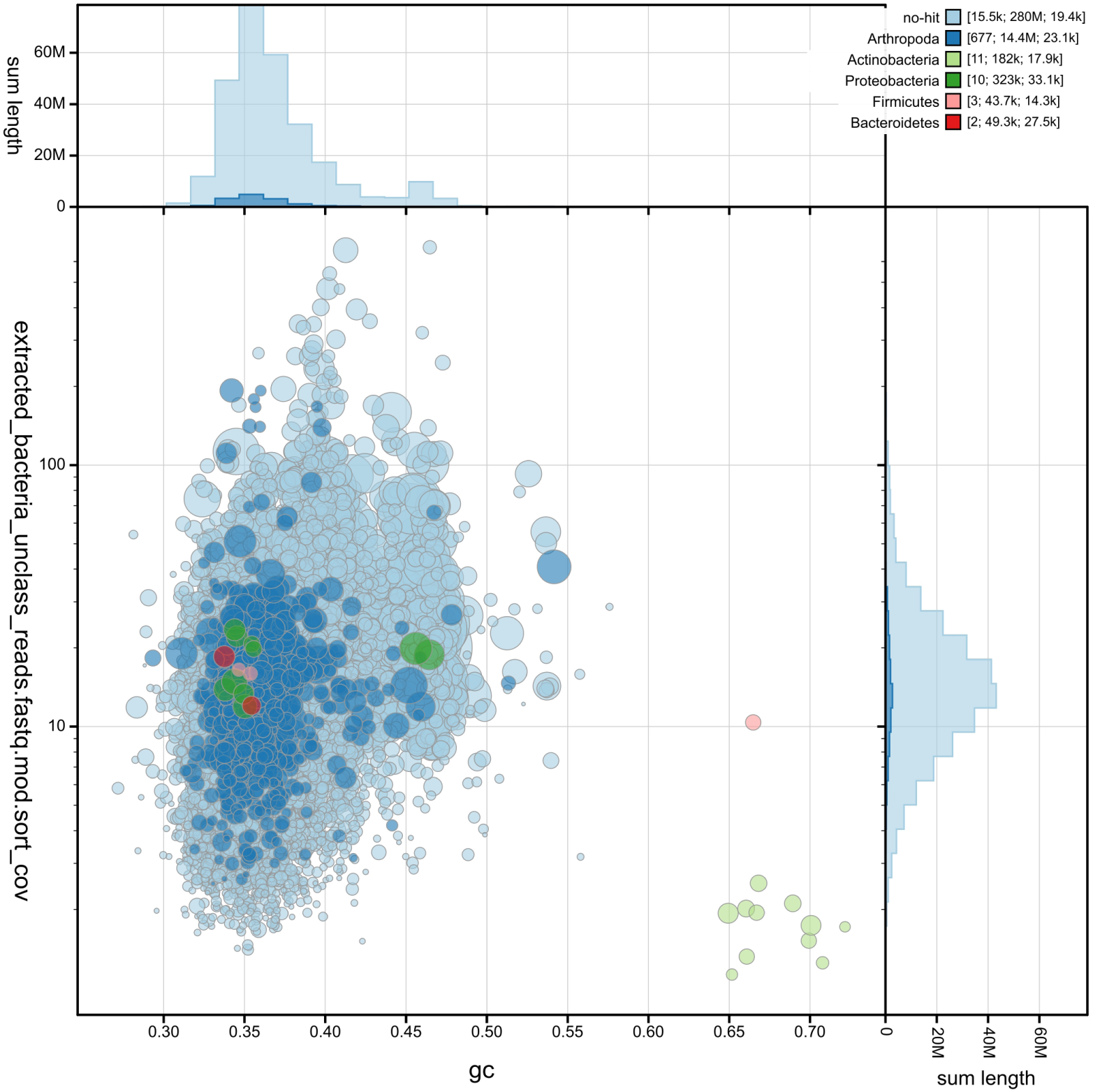
BlobToolKit circle plot of the bacterial scaffold fraction (16,216 contigs, 295 Mb). Arthropoda (677, 14.4 Mb) = beetle DNA; Actinobacteria (11), Proteobacteria (10), Firmicutes (3), Bacteroidetes (2) = genuine microbial associates; no-hit (15,534, 280 Mb). Despite 34 *Pseudomonas* contigs, tBLASTn for ndmA/B/C/D returned zero hits.

**Figure S2.**
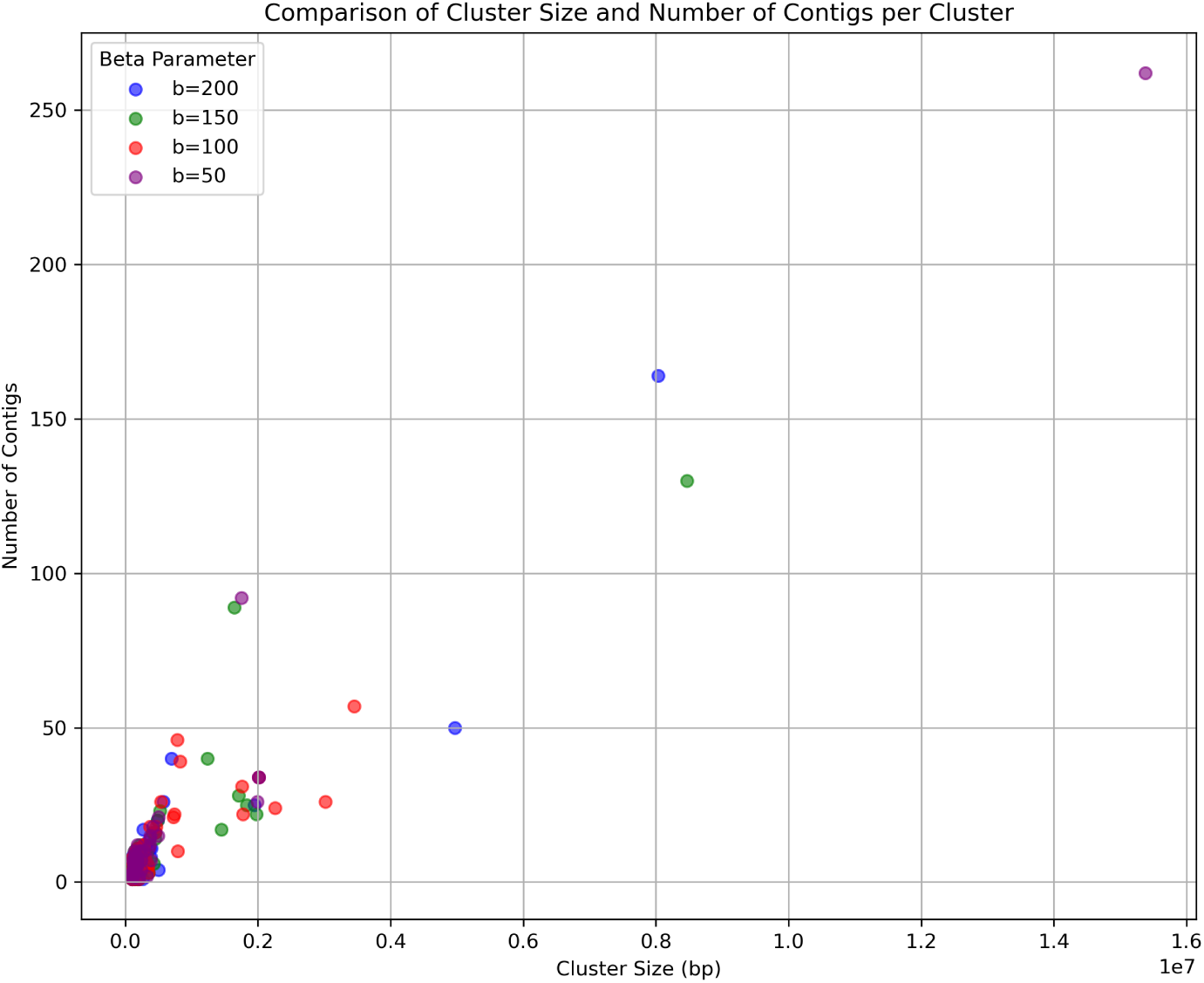
VAMB metagenomic binning parameter comparison. Cluster size vs number of contigs for beta = 50, 100, 150, 200, showing b = 50 produced finest-grained bins.

**Figure S3.**
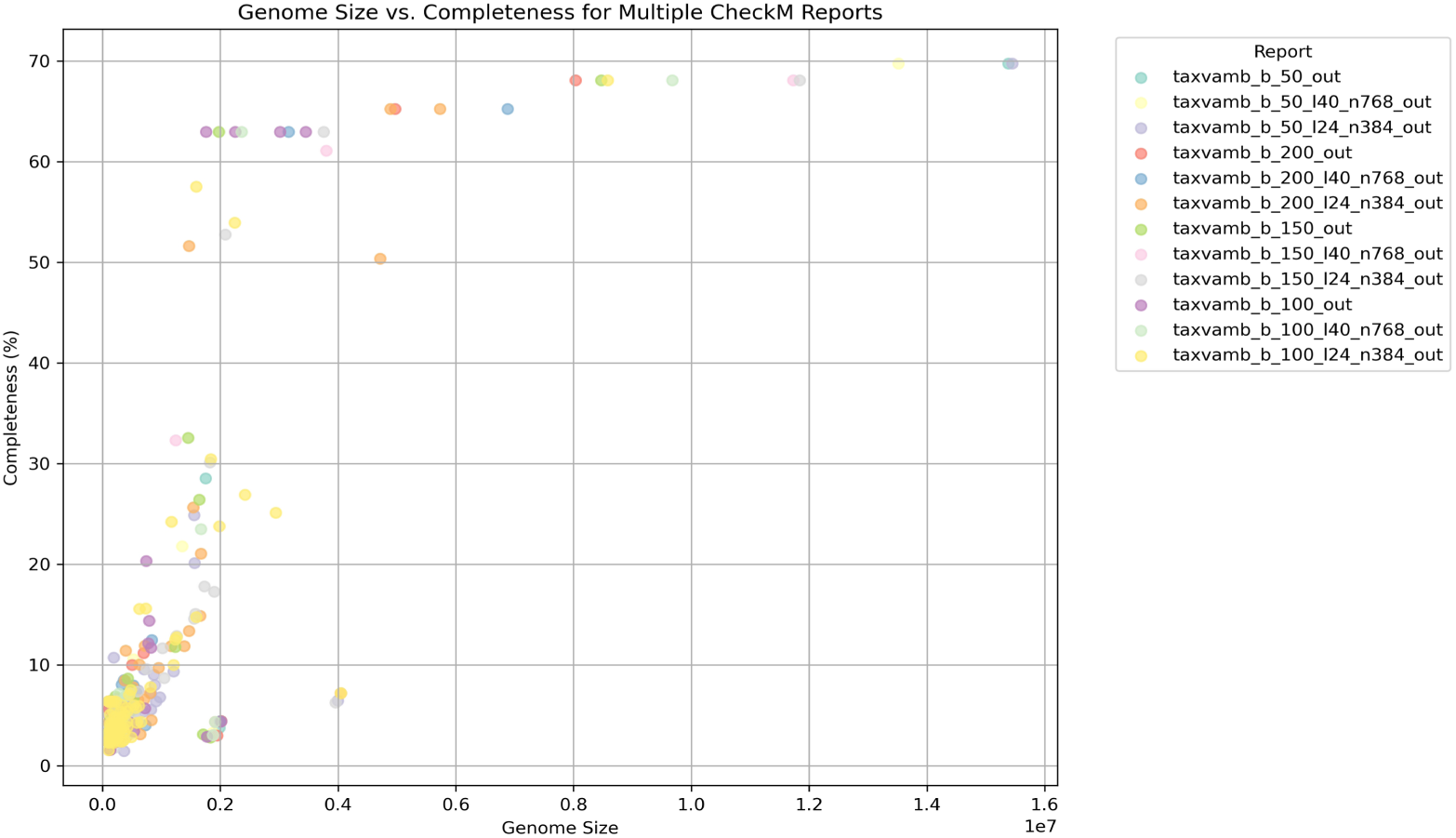
Genome size vs completeness scatter for all 12 VAMB runs. b = 200 produced largest bins (8-15 Mb, 65-70% complete) but with mixed content; b = 50 gave clean small bins.

**Figure S4.**
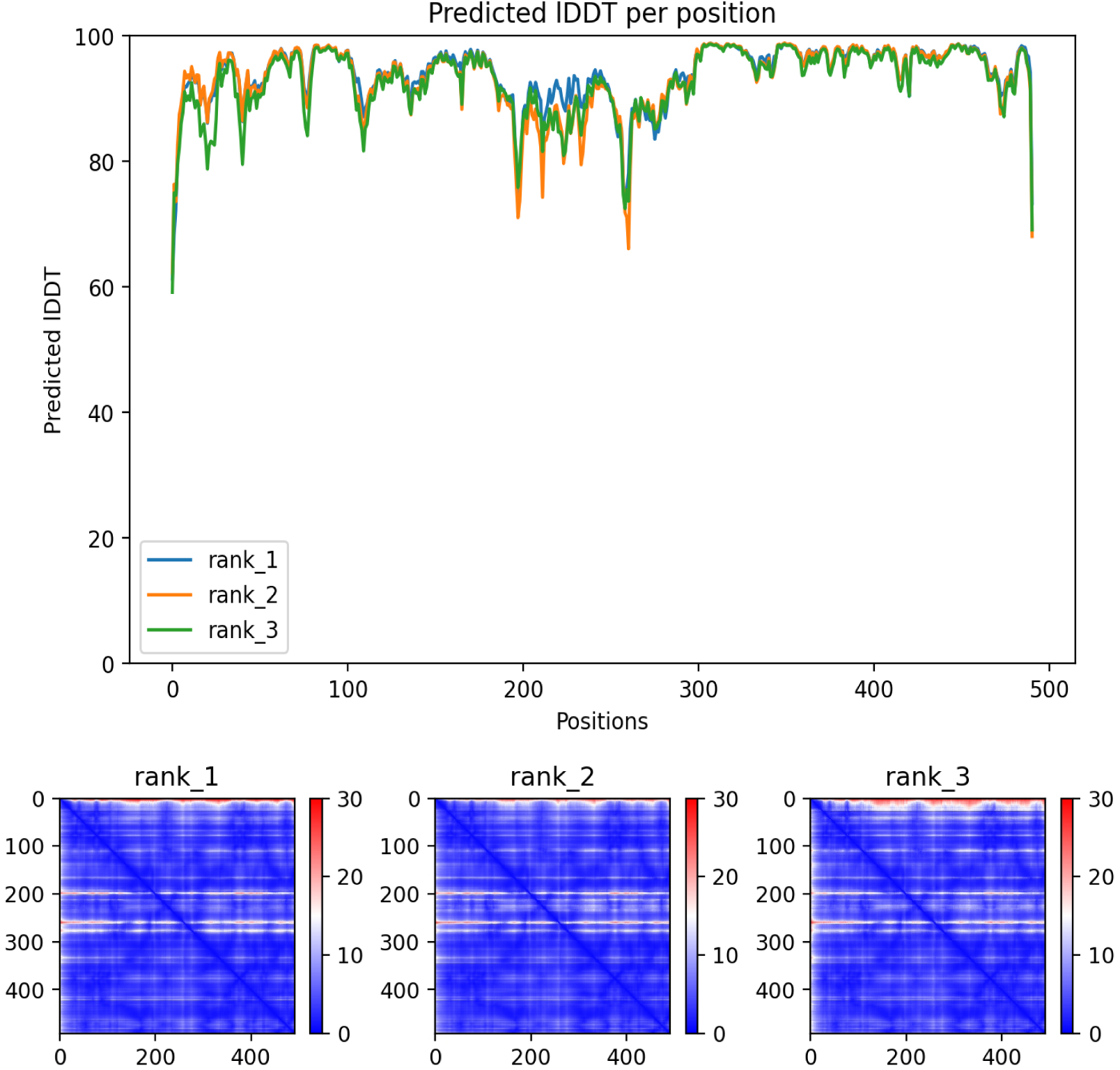
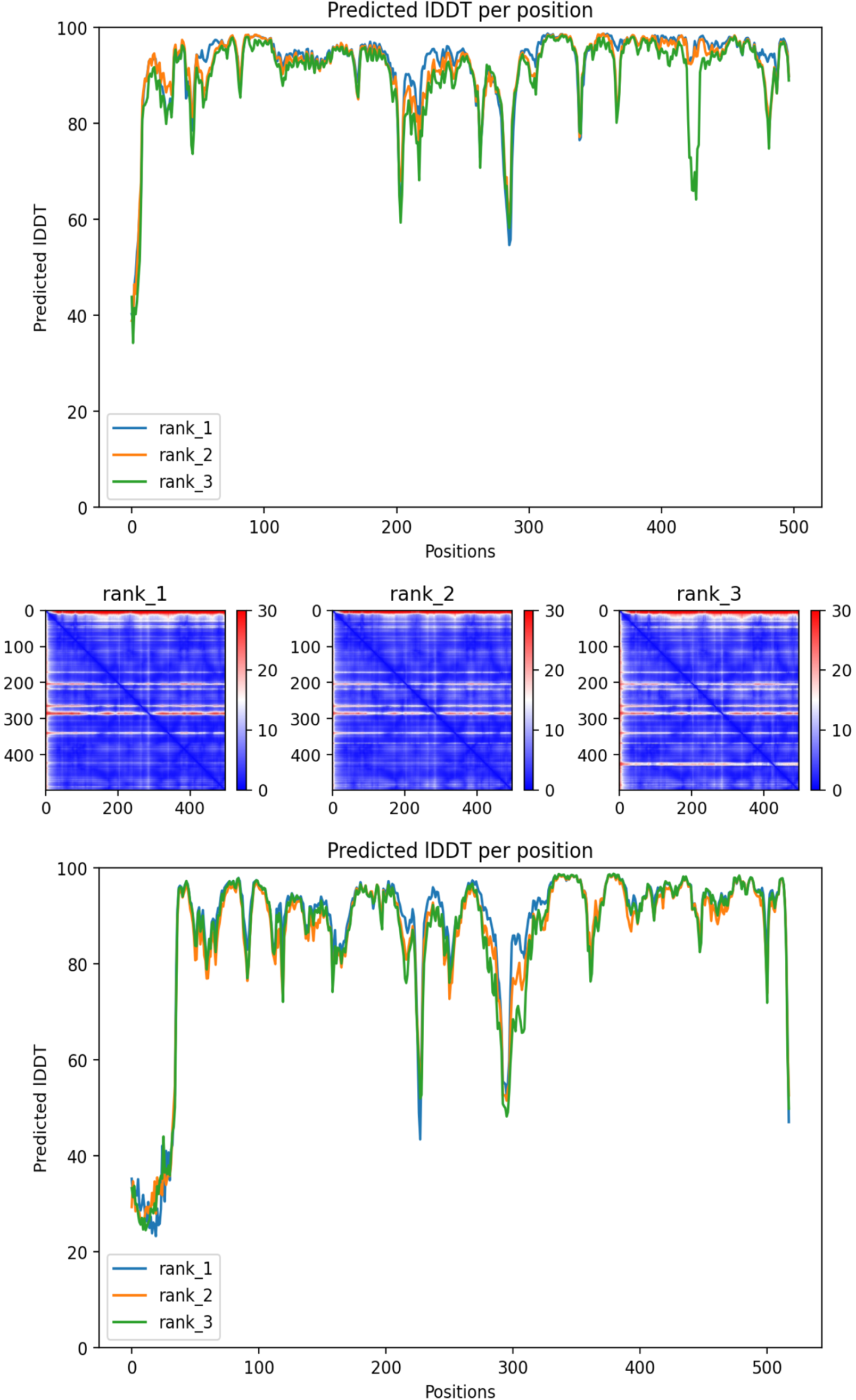
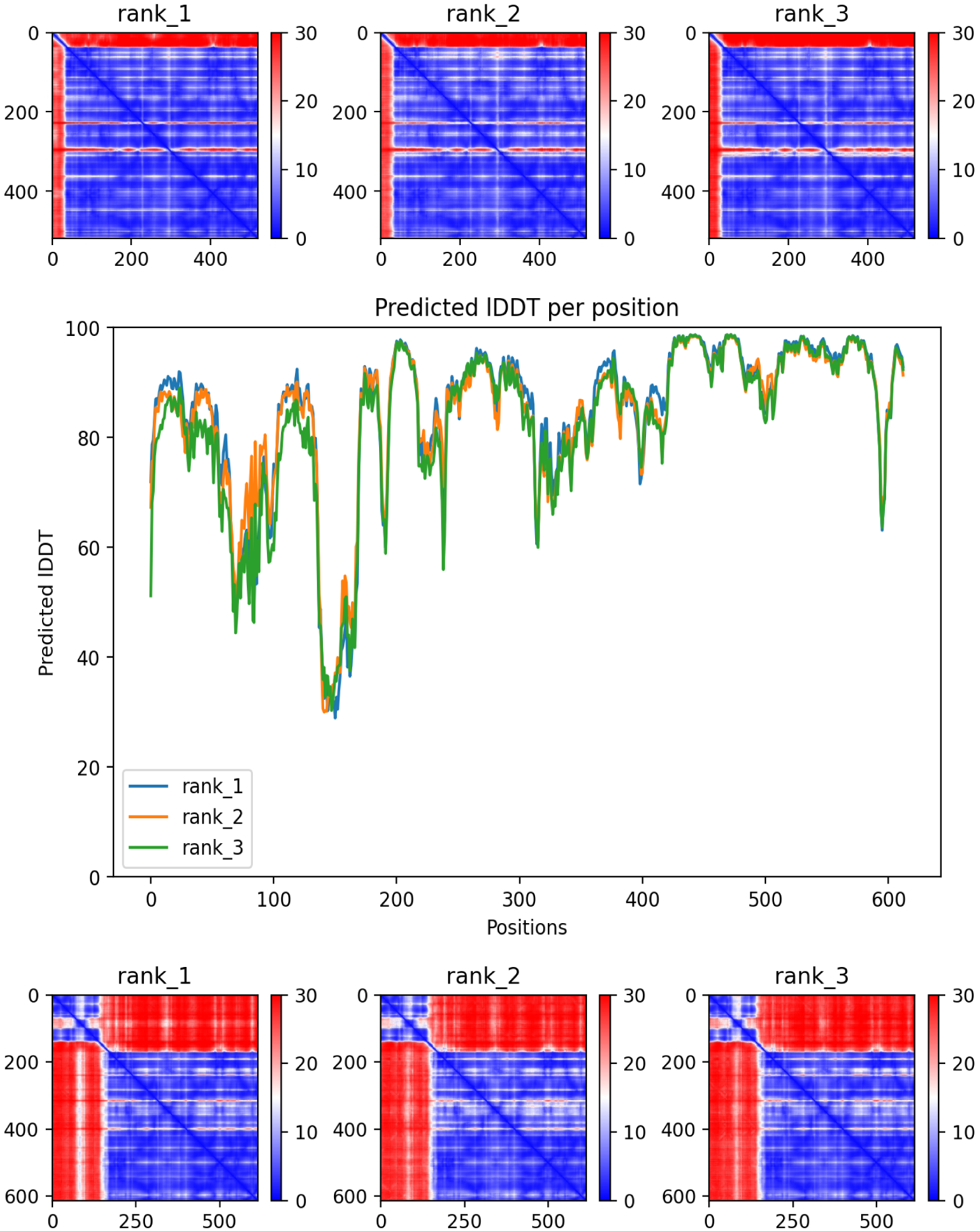
AlphaFold2 pLDDT and PAE confidence plots for all *Araecerus fasciculatus* P450 candidates (g3401, g9600, g17057, g11033).

**Data available at:** https://zenodo.org/records/20639466)

**Data S1.** TaxVAMB Taxometer classification results for all 16,216 bacterial scaffold contigs (results_taxometer.csv).

**Data S2.** VAMB cluster metadata for b = 50 bins (vaevae_clusters_metadata.tsv).

**Data S3.** CheckM2 quality report for 100 kb+ bins (quality_report.tsv).

**Data S4.** All supplementary data files are available from Zenodo (DOI: 10.5281/zenodo.20639466).

